# Ensembles and engrams in mouse cortical and sub-thalamic brain regions supporting context and memory recall

**DOI:** 10.1101/2025.06.29.662192

**Authors:** William W Taylor, Vienna Gao, Laura Korobkova, Brian G Dias

## Abstract

Associative learning supports learning about outcomes associated with contexts and cues. During learning, cellular ensembles that become active can be incorporated into a memory engram and later reactivated to support memory recall. Studies exploring engram formation and reactivation have primarily used contextual conditioning in mice and made little distinction between engrams encoding contextual information versus cue-associated learning and recall. Furthermore, often missing in such analyses is exploration of sex differences in engram profiles. Using auditory fear conditioning and activity-dependent tagging in mice, we set out to disaggregate context-associated engrams from those associated with learning and recall while also profiling potential sex differences. Specifically, we quantified cellular activity during context exposure, fear recall, extinction training, and extinction recall in cortical and subthalamic brain regions supporting learning and memory. We found that male mice had larger ensembles of cells active in the infralimbic prefrontal cortex (IL-PFC) during context exposure while female mice recalling a fear memory had a significantly greater proportion of cells that were active in the IL-PFC independent of context. Across sexes, we found greater reactivation of extinction engrams in the IL-PFC compared to contextual engrams. While we found ensembles and engrams in the prelimbic prefrontal cortex (PL-PFC) and zona incerta (ZI), no sex differences were noted in these regions. These results not only emphasize that there is a distinction to be made between ensembles and engrams encoding contextual information from those encoding cue-associated learning and recall, but also highlight sex differences in ensemble and engram allocation.

## INTRODUCTION

Associating contexts and environmental cues with outcomes and then later recalling these associations is integral to survival and our daily lives. Recently, we have come to appreciate that groups of neurons (ensembles) fire during associative learning with a subset of these neurons forming a memory trace (engrams) that is later reactivated to support memory recall (1–4). Further support for the role of such ensembles and memory engrams in successful learning and recall come from studies labeling cells in specific contexts or when specific cues are becoming associated with aversive or appetitive outcomes and later modulating their activity when recall of the relevance of contexts or cues is being tested. Broadly speaking, silencing cells that had been previously incorporated into engrams significantly disrupts memory recall, while activation can elicit expression of learned behaviors independent of any conditioned context or cue that had been encountered at the time of learning (4–12). Together such studies have established engrams as important units of memory in the brain that come to encode learning and support memory recall.

To our knowledge, studies using associative learning protocols have thus far made little to no distinction between ensembles encoding contextual information vs ensembles active at the time of learning and memory recall. Additionally, sex differences in ensemble size and engram reactivation patterns after associative learning and memory recall are not well understood. Filling these gaps in knowledge is important to understand not only disruptions to context- and cue-related learning and memory recall, but also sex differences in these disruptions that are reported in anxiety and trauma-related conditions like Post Traumatic Stress Disorder (PTSD) (13–15). To eventually achieve these objectives, in this study, we trained TRAP2 (**T**argeted **R**ecombination in **A**ctive **P**opulations); floxed-tdTomato male and female mice to associate conditioned stimulus (CS+) tone presentations in Context A with mild footshocks. Next, we activated the TRAP2 system (16) to specifically label cells during one of two conditions: either during exposure to Context B alone without any extinction training (context group), or alternatively, during extinction training when animals were exposed to CS+ tone presentations without footshocks in Context B (extinction group). This genetic tagging approach allowed us to capture and distinguish cellular ensembles activated under these experimental conditions. Finally, we stained for C-FOS following CS+ presentations without footshocks in Context B, which constituted a fear recall test in the context group and an extinction recall test in the extinction group. This approach allowed us to investigate ensemble size and engram reactivation patterns during context exposure, extinction training, fear recall, and extinction recall in cortical and sub-thalamic brain regions involved in learning and memory: the prelimbic prefrontal cortex (PL-PFC), infralimbic prefrontal cortex (IL-PFC), and zona incerta (ZI). Briefly, our results demonstrate sex differences, specifically in the IL-PFC, in the size of ensembles active during context exposure and the reactivation of engrams during fear recall, as well as preferential IL-PFC reactivation during extinction recall in both sexes.

## METHODS

### Animals

TRAP2 mice (Fos^tm2.1(icre/ERT2)Luo^/J, strain 030323) and floxed-tdTomato reporter mice (B6.Cg-Gt(ROSA)26Sor^tm14(CAG-tdTomato)Hze^/J, strain 007914) were obtained from Jackson Laboratory (Bar Harbor, ME, USA) and crossed to generate adult male and female TRAP2/+;floxed-tdTomato/+ mice for all experiments (n = 17 females, 14 males). These mice allow for fluorescent tdTomato expression in CRE-expressing cells when 4-Hydroxytamoxifen (4OHT) is present (Supplemental Figure 1) (16). Genotyping was performed using Jackson Laboratory “Quick DNA purification protocol” from ear clips. The mice were group housed and maintained on *ad libitum* food and water on a 12-hour light/dark cycle. Both male and female mice were included in all experiments. Experiments were approved by the Institutional Animal Care and Use Committee of Children’s Hospital Los Angeles and followed NIH standards.

### Auditory fear conditioning, context exposure, extinction training, and extinction recall

All behavioral sessions were conducted in conditioning chambers (Coulbourn Instruments) which included tone generators (internal speakers) and shock generators (cage-floor inserts) controlled by FreezeFrame software (Actimetrics). Freezing behavior was recorded and measured via video algorithms embedded in FreezeFrame software. All behavioral sessions took place during the light cycle and animals were returned to the vivarium afterwards.

*Day 0: Habituation to Context A.* Mice were habituated to Context A, which consisted of metal rod flooring, chamber and room lights off, Peroxigard as a cleaning agent, and an infrared light to allow for recording.

*Day 1: Fear conditioning in Context A.* The mice underwent auditory fear conditioning, consisting of a 180-second baseline period followed by five 30-second 6 kHz tone (T) presentations, the conditioned stimulus (CS+), each co-terminating with a one-second foot shock (0.5 mA), the unconditioned stimulus (US). Tone-shock pairing were presented with 30- second inter-trial intervals (ITI).

*Day 2: Habituation to Context B.* Mice were habituated to Context B, which was the same behavioral chamber but with distinct contextual cues including a plexiglass floor, chamber and room lights on, and 70% ethanol as a cleaning agent.

*Day 3: 4OHT injection and extinction learning or context exposure in Context B.* One hour before extinction training or context exposure, all mice were injected with 4OHT. 4OHT was generously provided by the NIMH Chemical Synthesis and Drug Supply Program or purchased from Sigma-Aldrich (St. Louis, MO,USA; catalogue #H6278). 4OHT was prepared fresh on the day of injection, as described by DeNardo et al., 2019. Briefly, 4OHT was dissolved in ethanol at 20 mg/ml by shaking at 37°C. Corn oil (Sigma-Aldrich, catalogue #C8267) was added for a final concentration of 10mg/ml and ethanol was evaporated. Mice were injected intraperitoneally (i.p.) at a dose of 50 mg/kg. The extinction group (n = 18) received a 180-second baseline period followed by 30 CS+ tone presentations, separated by 30-second ITIs, without any footshock to facilitate extinction learning. The context group (n = 13) was placed in Context B without being exposed to any tones for the same amount of time as the extinction training session lasted. Animals were returned to the vivarium and left undisturbed for 4 days to allow for robust expression of tdTomato in TRAPped cells.

*Day 8: Extinction recall or fear recall testing in Context B and brain collection.* All mice were returned to Context B and exposed to 5 CS+ presentations without foot shocks, separated by 30-second ITIs. For the extinction group, this protocol tested the recall of extinction learning and in the context group this tested the recall of the initial fear conditioning. Mice were deeply anesthetized with a mixture of ketamine and dexdomitor, 60-90 minutes after testing for recall, and transcardially perfused with ice-cold phosphate-buffered saline (PBS), followed by 4% paraformaldehyde (PFA) in PBS, and brains were collected for further analysis.

### Immunohistochemistry

Brains were fixed overnight in 4% PFA in PBS and then transferred to a 30% sucrose solution in PBS for 3-4 days until fully saturated. Once saturated, brains were flash-frozen and coronally sectioned at 35 µm using an Epredia CryoStar NX70. Sections were mounted onto slides and stored at -80°C without light exposure. For C-FOS immunohistochemistry, brain sections were washed in 1X PBS and blocked for one hour with 5% normal goat serum and 0.1% Triton-X. Sections were incubated overnight at 4°C with Rabbit anti C-FOS (1:500; sc-52, Santa Cruz Biotechnology, Dallas, TX). Sections were washed then incubated for 2 hours at room temperature with Goat anti-Rabbit Alexa Fluor™ 488 (1:1,000; A11034, ThermoFisher, Waltham, MA). Nuclei were counterstained with Hoechst (1:10,000; H3570, Invitrogen, Waltham, MA). Images were acquired using a Leica STELLARIS 5 WLL microscope with a 20x/0.75 objective.

### Image and data analysis

Image analysis was conducted using QuPath software (17). Regions of interest were identified and delineated based on the Allen Mouse Brain Atlas (mouse.brain-map.org). TRAP+ and C-FOS+ cells were automatically detected using a consistent threshold and manually confirmed for accuracy. Raw cell counts were measured for each region by analyzing 2-3 images per animal and the average density of TRAP+, C-FOS+ and TRAP+/C-FOS+ cells (cells per mm²) was calculated with individual animals serving as the unit of analysis. To compute the proportion of newly activated cells, the number of exclusively C-FOS+ (TRAP-) cells was divided by total C-FOS+ cells. To compute percentages of reactivated cells, the number of double labelled cells (TRAP+ and C-FOS+) was divided by total TRAP+. Statistical analyses for both image analysis and behavior were conducted using RStudio (version 4.3.2). Individual group or sex comparisons were done using the Wilcoxon signed-rank test and posthoc testing was conducted using the same and Tukey HSD.

## RESULTS

### Male and female mice show similar freezing behavior during context exposure and fear recall

The context group initially underwent fear conditioning consisting of 5 CS+ (tone) paired with unconditioned stimulus (US, footshock) in Context A (Figure 1, Day 1). A repeated measures ANOVA was conducted to examine the effects of sex and tone presentation on freezing behavior. There was no significant main effect of sex (F(1, 11) = 0.088, p > 0.05). However, there was a significant main effect of tone presentation (F(4, 44) = 32.785, p < 0.001). Post-hoc testing revealed that both male and female mice froze significantly more to the 5^th^ tone presentation than the 1^st^, indicating they successfully formed an association between the CS+ and US (both p < 0.0001). There was also a significant interaction between sex and tone presentation (F(4, 44) = 3.492, p = 0.01), but post-hoc testing comparing freezing in males and females at each CS+ presentation revealed no significant differences at any specific tone presentation. Following fear conditioning, both males and females demonstrated similar levels of freezing when exposed to Context B (W = 10, p > 0.05) (Figure 1, Day 3) after being injected with 4OHT 60 minutes prior to context exposure. Finally, for fear memory recall, animals were exposed to 5 tone presentations in Context B (Figure 1, Day 9). Males and females did not differ in their freezing behavior during the 5 tone presentations (W = 19, p > 0.05). A repeated measures ANOVA comparing context exposure with fear recall sessions revealed a significant main effect of session (F(1, 11) = 33.06, p < 0.001) and no main effect of sex (F(1, 11) = 0.533, p > 0.05) or any significant interaction between sex and session (F(1, 11) = 1.218, p > 0.05). Post-hoc testing collapsed by sex showed that mice froze significantly more during fear recall than context exposure (V = 1, p < 0.001). For full behavioral data see Supplemental Figure 2. Comparing freezing during context exposure and fear recall revealed no significant effect of sex (F(1, 11) = 0.533, p > 0.05) or interaction of sex and session interaction F(1, 11) = 1.218, p > 0.05), but a significant effect of session (F(1, 11) = 33.06, p < 0.001) and post-hoc testing collapsed by sex showed that mice froze significantly more during fear recall than context exposure (V = 1, p < 0.001). Data show percent time freezing and are represented as mean±SEM.

**Figure 1.**
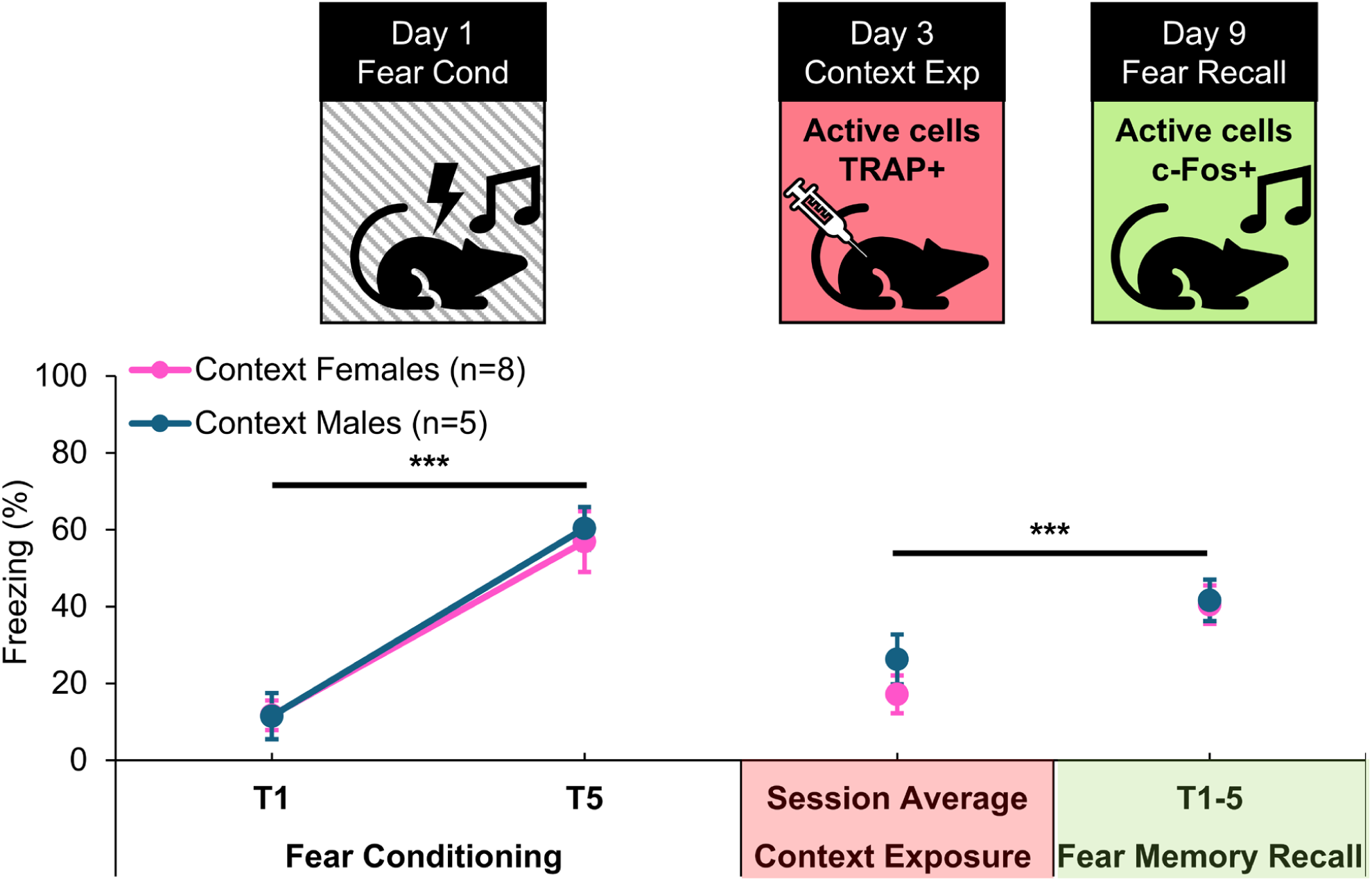
Fear behavior during fear conditioning, contextual exposure, and fear memory recall did not differ by sex. Both male and female mice in the context exposure group significantly increased their freezing in response to the CS+ over the course of fear conditioning, showing greater freezing to the 5^th^ tone presentation compared with the 1^st^ (both p <0.0001). There were no sex differences in freezing to a neutral context during context exposure. There were also no sex differences in freezing to 5 tone presentations during fear recall.

### Sexually dimorphic ensemble activation during context exposure in IL-PFC, but not PL- PFC and ZI

Density of cellular ensembles active during context exposure in the PL-PFC, IL-PFC, and ZI were compared between the sexes (Figure 2A-C). Compared to female mice, male mice had significantly larger ensembles of cells labeled in the IL-PFC (W = 36, p = 0.02), but not in the PL-PFC or the ZI (both p > 0.05) upon exposure to Context B (Top Row). No significant differences in the density of ensembles active during fear recall testing in Context B (Figure 2D- F) were found in the PL-PFC, IL-PFC, or ZI between male and female mice (all p > 0.05) (Middle Row). Similarly, we found no sex differences in the density of engram cells that were active during Context B exposure and also active during fear recall testing in Context B in the PL-PFC, IL-PFC, and ZI (all p > 0.05) (Figure 2G-I) (Bottom Row).

**Figure 2.**
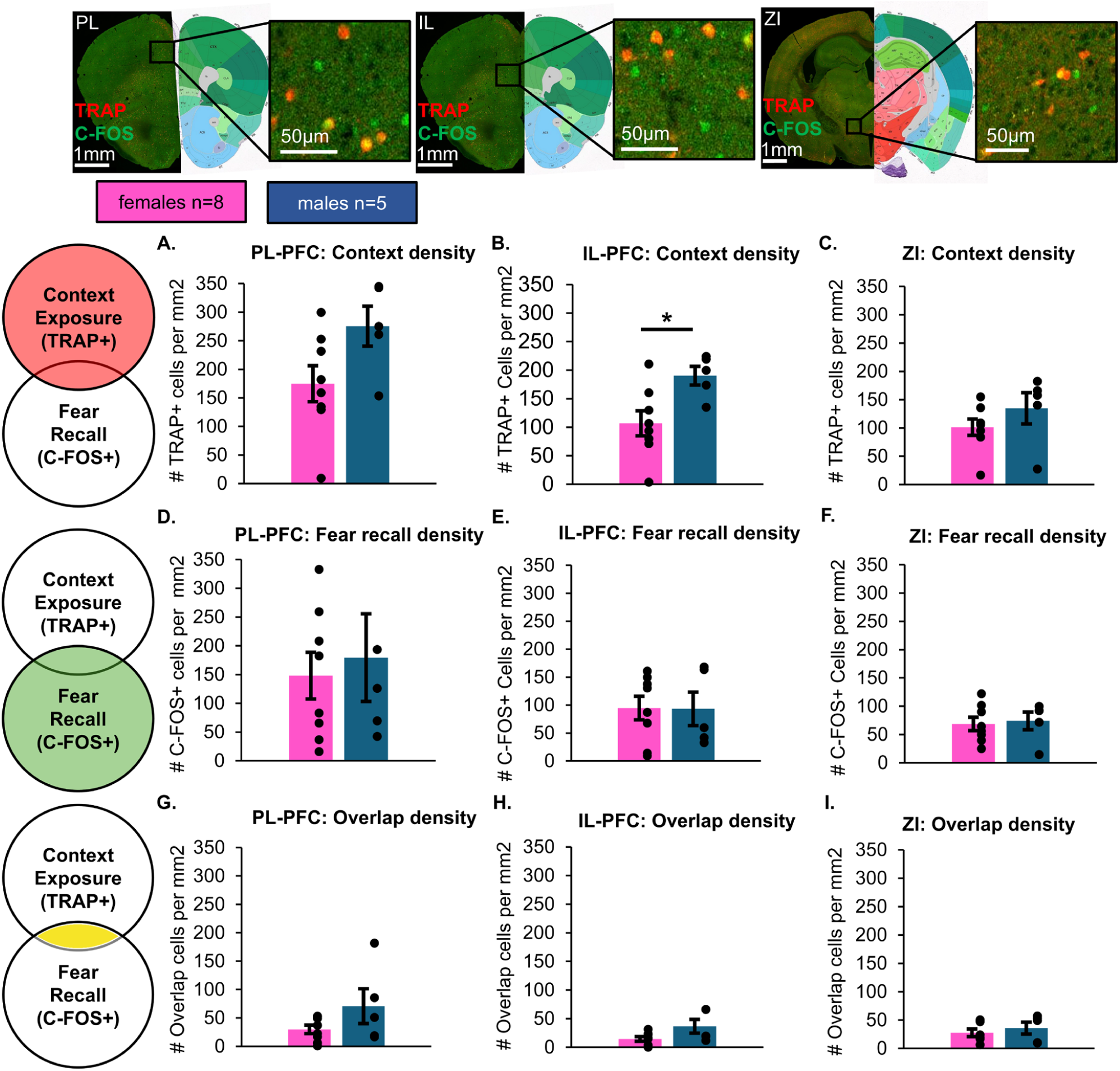
Sex differences in ensemble activation during context exposure in IL-PFC, but not PL-PFC and ZI. **(A-C)** Density of ensemble activation during Context B exposure was measured in the PL-PFC, IL-PFC, and ZI. Male mice had significantly larger ensembles active in the IL-PFC during context exposure (W = 36, p = 0.02). Data show mean # of TRAP+ cells per mm^2^±SEM. **(D-F)** Density of ensemble activation during fear recall in Context B was measured in the PL-PFC, IL-PFC, and ZI, showing no significant sex differences. Data show mean # of C-FOS+ cells per mm^2^±SEM. **(G-I)** Density of engram cells active both during context exposure and fear recall was measured in the PL-PFC, IL-PFC, and ZI, showing no significant sex differences. Data show mean # of both TRAP+ and C-FOS+ cells per mm^2^±SEM.

### Male and female mice show similar freezing behavior during extinction learning and recall

Similarly to the context group, the extinction group initially underwent fear conditioning in Context B (Figure 3, Day 1). A repeated measures ANOVA revealed no main effect of sex (F(1, 16) = 0.474, p > 0.05), but a main effect of tone presentation (F(4, 64) = 31.021, p < 0.001) and no significant interaction (F(4, 64) = 1.549, p > 0.05). Post-hoc testing revealed that by the 5^th^ tone presentation of fear conditioning, both male and female mice of the extinction group successfully associated the tone with the footshock and demonstrated significantly more freezing behavior to the 5^th^ tone than initially observed to the 1^st^ tone presentation (both p < 0.0001). During extinction training (Figure 3, Day 3), animals were exposed to 30 tone presentations in the absence of footshock in Context B, after being injected with 4OHT 60 minutes prior to extinction training. A repeated measures ANOVA revealed no main effect of sex (F(1, 11) = 0.533, p > 0.05), but that a main effect of CS+ bin (5 tone bins) was significant (F(1, 11) = 33.060, p < 0.001). This indicates that freezing behavior varied significantly across time as animals underwent extinction training, but did not differ by sex. There was no significant interaction between sex and CS+ bin (F(1, 11) = 1.218, p > 0.05). Post-hoc testing collapsed by sex showed that mice froze significantly more to the first bin of 5 tone presentations than the last bin (V = 162, p < 0.001).

**Figure 3.**
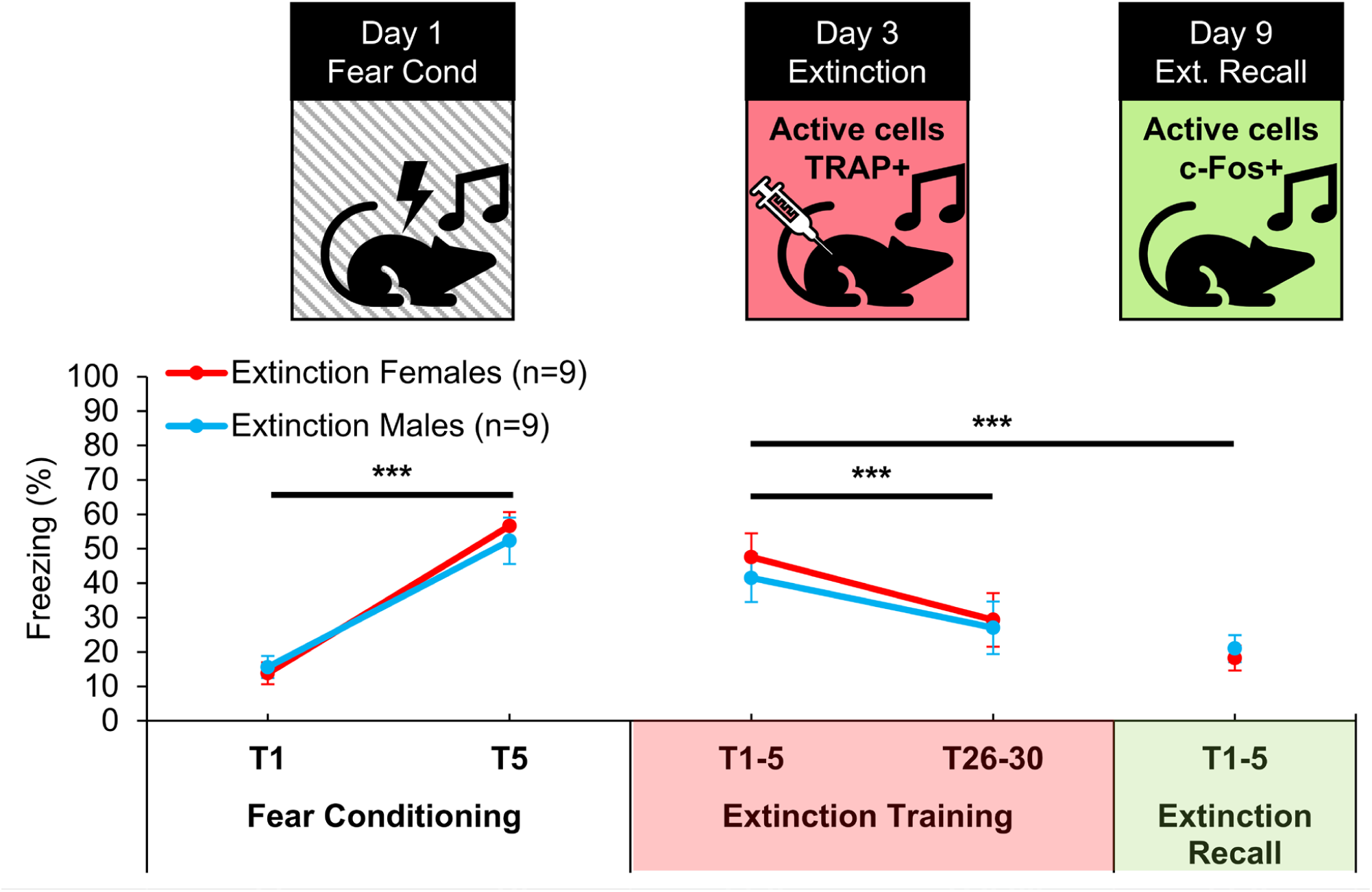
Fear behavior during fear conditioning, extinction training, and extinction memory recall did not differ by sex. Both male and female mice in the extinction group significantly increased their freezing in response to the CS+ over the course of fear conditioning in Context A, showing greater freezing to the 5^th^ tone presentation compared with the 1^st^ (both p < 0.0001). There were no sex differences in freezing during extinction training in Context B. Comparing freezing during 5 tone bins of extinction training by sex revealed no significant effect of sex (F(1, 11) = 0.533, p > 0.05), a significant main effect of CS+ bin (F(1, 11) = 33.060, p < 0.001), and no significant interaction of sex and tone bin (F(1, 11) = 1.218, p > 0.05). Post-hoc testing collapsed by sex showed that mice froze significantly more during the first 5 tone bin than the last (V = 162, p < 0.001). There were no sex differences in freezing during extinction recall in Context B (W = 47, p > 0.05). Comparing the first 5 tone bin of extinction training (fear recall) with extinction recall revealed a significant main effect of session (F(1, 16) = 58.788, p < 0.0001), no effect of sex (F(1, 16) = 0.052, p > 0.05), and no interaction between session and sex (F(1, 16) = 1.842, p >0.05). Post-hoc testing collapsed by sex showed that mice froze significantly more during the first 5 tone presentations of extinction training than during extinction recall (V = 171, p < 0.0001). Data show percent time freezing and are represented as mean±SEM.

There were no sex differences in freezing during extinction recall in Context B (Figure 3, Day 9) (W = 47, p > 0.05) and both male and female mice showed successful extinction recall, with a repeated measures ANOVA comparing the first bin of extinction training (fear recall) with extinction recall revealing a significant main effect of session (F(1, 16) = 58.788, p < 0.0001), no effect of sex (F(1, 16) = 0.052, p > 0.05), and no effect of the interaction between session and sex (F(1, 16) = 1.842, p > 0.05). Post-hoc testing collapsed by sex showed that mice froze significantly more during fear recall than during extinction recall (V = 171, p < 0.0001). For full behavioral data, including extinction curves, see Supplemental Figure 3.

### Male and female mice show similar ensemble activation during extinction training and recall

There was no difference between the density of cellular ensembles active during extinction training (Figure 4A-C) in the PL-PFC, IL-PFC, or in the ZI of male and female mice (all p > 0.05) (Top Row). Similarly, there were no significant sex differences in the density of cellular ensembles active during extinction recall (Figure 4D-F) in the PL-PFC, IL-PFC, or in the ZI (all p > 0.05) (Middle Row). There were also no sex differences in the density of overlapping engram cells in the PL-PFC, IL-PFC, and ZI that were active both during extinction training and active during extinction recall (Figure 4G-I) (all p > 0.05) (Bottom Row).

**Figure 4.**
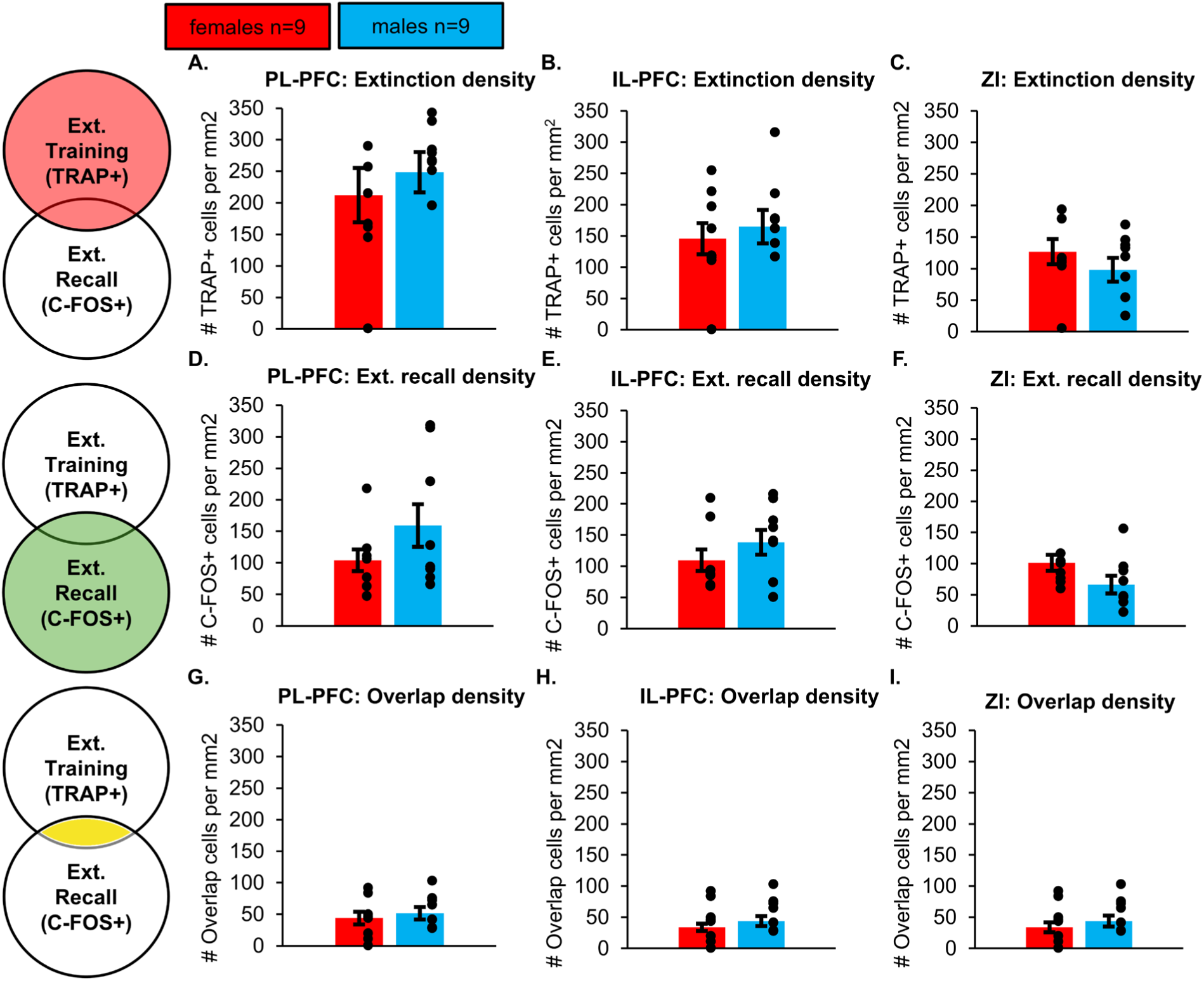
No sex differences in ensemble activation during extinction training, extinction recall, or engram reactivation during extinction recall. (A-C) Density of ensemble activation during extinction training was measured in the PL-PFC, IL-PFC, and ZI and no significant sex differences were found. Data show mean # of TRAP+ cells per mm^2^±SEM. **(D-F)** Density of ensemble activation during extinction recall was measured in the PL-PFC, IL-PFC, and ZI and no significant sex differences were found. Data show mean # of C-FOS+ cells per mm^2^±SEM. **(G-I)** Density of engram cells active both during extinction training and extinction recall was measured in the PL-PFC, IL-PFC, and ZI and and no significant sex differences were found. Data show mean # of both TRAP+ and C-FOS+ cells per mm^2^±SEM.

### Sexually dimorphic activation in the IL-PFC during fear recall and preferential reactivation of IL-PFC during extinction recall

The percentage of all cells active during fear recall or extinction recall (C-FOS+) that were newly active (C-FOS+ and TRAP-) was analyzed as a function of group and sex (Figure 5A-C). A two-way ANOVA revealed no main effect of group (p > 0.05), but a significant main effect of sex (F(1, 27) = 4.66, p = 0.04) and a significant interaction between group and sex (F(1, 27) = 7.71, p < 0.01). Tukey post-hoc testing revealed that female mice exposed to Context B had significantly more newly active cells than male mice that had been exposed to Context B (p = 0.01) or females that underwent extinction training (p = 0.046). There was no main effect of, or interaction between, group and sex in the PL-PFC and ZI (all p > 0.05).

**Figure 5.**
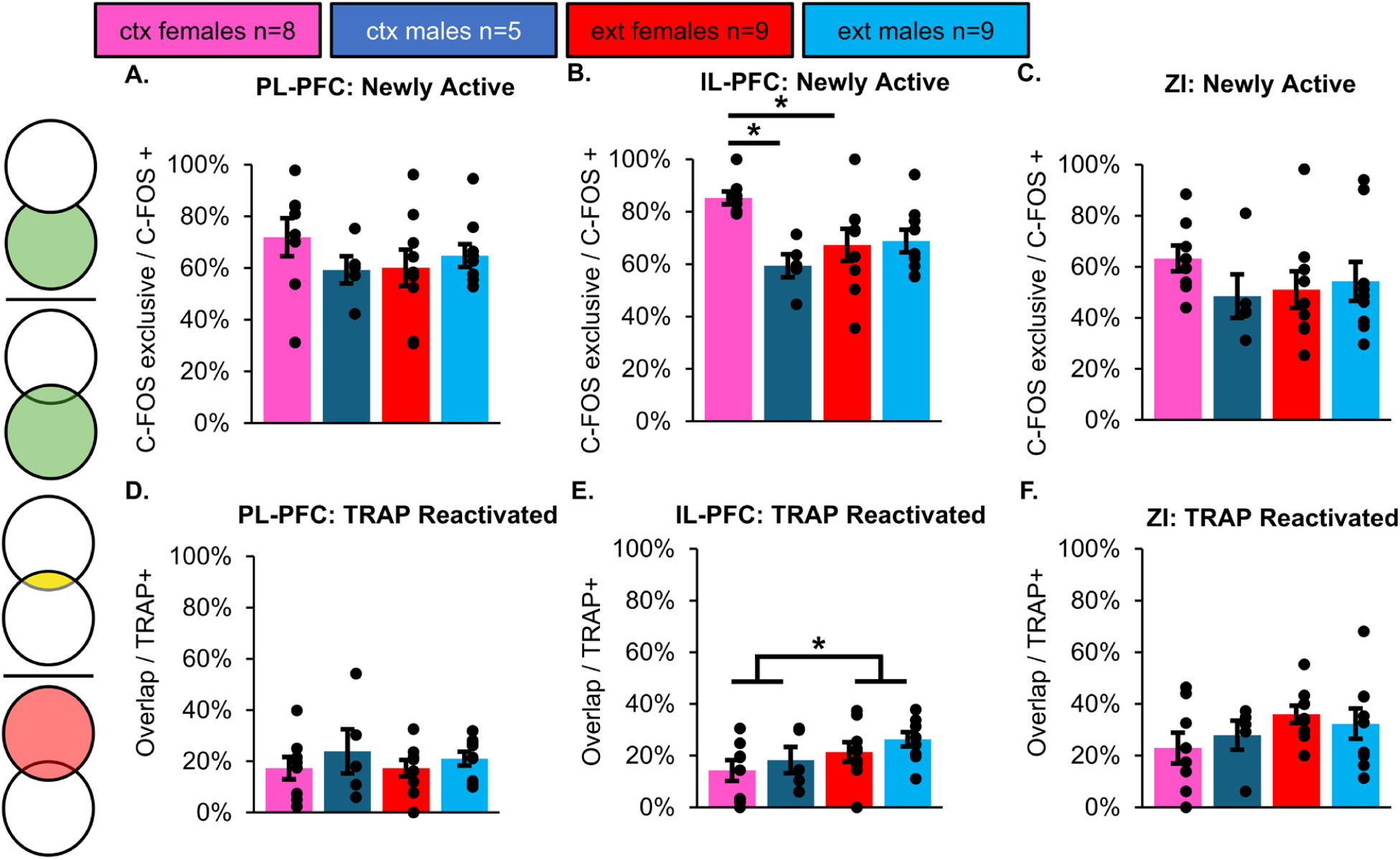
Sex differences in patterns of ensemble and engram activation during recall in IL-PFC, but not PL-PFC and ZI. (A-C) The percentage of newly active cells (exclusively C-FOS+ divided by all C-FOS+) was analyzed as a function of group and sex. Comparing % newly active by group and sex revealed no main effect of group (F(1, 27) = 1.58, p > 0.05), but a significant main effect of sex (F(1, 27) = 4.66, p = 0.04) and a significant interaction between group and sex (F(1, 27) = 7.71, p < 0.01). Posthoc testing revealed that female context exposed mice had a greater % of newly active cells compared with male context exposed mice (p = 0.01) or females that underwent extinction training (p = 0.046). There was no main effect of, or interaction between, group and sex in the PL-PFC and ZI (all p > 0.05). Data show mean % newly active±SEM. **(D-F)** The percentage of reactivated cells (double labelled cells divided by all TRAP+ cells) was analyzed as a function of group and sex. Comparing % reactivation by group and sex revealed a trending main effect of group in the IL-PFC (F(1, 27) = 3.82, p = 0.06), no main effect of sex (F(1, 27) = 2.038, p > 0.05), and no interaction between group and sex (F(1, 27) = 0.014, p < 0.05). Collapsing across sex revealed that the extinction group had a significantly higher % reactivation than the context exposure group (W = 167.5, p = 0.045) (Supplemental Figure 4). Data show mean % reactivation±SEM.

The percentage of all cells labelled during context exposure or extinction learning (TRAP+) that were reactivated (TRAP+ and C-FOS+) was analyzed as a function of group and sex (Figure 5D-F). A two-way ANOVA revealed a trending main effect of group in the IL-PFC (F(1, 27) = 3.82, p = 0.06), no main effect of sex (F(1, 27) = 2.038, p > 0.05), and no interaction between group and sex (F(1, 27) = 0.014, p < 0.05). Collapsing across sex and comparing engram reactivation in the IL-PFC between the groups revealed that there was significantly more reactivation in the extinction group than the context exposure group (W = 167.5, p = 0.045) (Supplemental Figure 4). There was no main effect of, or interaction between, group and sex in the PL-PFC and ZI (all p > 0.05).

## DISCUSSION

Ensembles of cells that are active during learning come to be incorporated into memory engrams. Engrams are distinct cellular representations of specific memories that may be reactivated to support future memory recall. To address potential sex differences in the allocation and reactivation of engrams across various stages of learning and memory, we used activity-dependent cell tagging using TRAP2; floxed-tdTomato mice to determine the density and reactivation patterns of engrams supporting context exposure, extinction training, fear recall, and extinction recall in male and female mice. Following auditory fear conditioning in Context A, we TRAPped cells active during extinction training or context exposure in the absence of extinction training, both, in Context B. We then stained for C-FOS following 5 tone presentations in Context B to identify active cells during extinction recall in the group that underwent extinction training and during fear recall in the group that underwent context exposure without extinction training. Given our laboratory’s interest in the influence of the ZI on learning and memory (18–20), the role of the PL- and IL-PFC in consolidation and recall of memory (21–23), and published literature from colleagues demonstrating the role of PFC→ZI communication in extinction learning and recall (24, 25), we chose to focus our attention on the PL-PFC, IL-PFC and the ZI. Our data suggest that there are sex differences in density of cells active in the IL-PFC during exposure to a context and in the percent of newly active cells during fear recall. Furthermore, we find enhanced reactivation of IL-PFC engrams during extinction recall, emphasizing the importance of this brain region in extinction learning and recall.

Sex differences in context- and cue-related learning and memory have been reported (13, 26). Efforts to model these sex differences preclinically in rodents have produced mixed results with some studies showing sex differences in fear- and extinction-related learning and memory, while others do not (14, 27). Here, we demonstrate no significant sex differences in behavior during any of the learning and recall tests after auditory fear conditioning in mice. Therefore, while our results demonstrate reliable fear acquisition to the CS+ during fear conditioning, successful within-session extinction learning, successful extinction recall, and high levels of fear recall in the context group, it might seem puzzling why we proceeded to count density of cells activated at these different epochs of behavior and reactivation profiles. First, these behavioral results provide confidence that our investigation of cellular activity successfully targeted cells active during the intended phase of learning and recall. Second, there are reports to suggest sex differences in neurobiology that are not directly linked to sex differences in behavior (28, 29). For example, a recent study used whole brain clearing to visualize TRAPped cells following recall of contextual fear and found that while male and female mice did not differ in their behavior, there was strong sexual dimorphism in the activation of different regions that correlated with fear behavior (30). Therefore, we posited that similar behavioral outputs in male and female mice in our experiments does not preclude the possibility of finding sex differences in density of cells active at the time of learning and sex differences in engram reactivation at various epochs of learning and memory. Our data suggest that while sex differences in density of cells active at the time of learning and sex differences in engram reactivation may exist, they could well occur independently of sex differences in fear behavior. This leaves open the possibility that the sex differences we observe in IL-PFC activation patterns could underly behaviors relevant to these tests that were not directly measured. For example, the PFC has been shown to support arousal, guide movement and action temporally, and provide higher-order processing of auditory information – all functions that would be employed during exploration of new environments and attending to auditory cues (31–34).

Much of our current understanding of memory engrams comes from literature focused on male mice and the use of contextual fear conditioning protocols. This pairing of context and fear does not allow for the dissociation of ensembles and engrams supporting fear- or extinction-related memory from those solely representing contextual information. Our experimental design allowed us to investigate sex differences in the activity of cellular ensembles during context exposure, independent of any CS+ presentations, and the reactivation of engrams in the same context during fear recall. We found that males have a denser ensemble of active cells in the IL- PFC during context exposure compared with females. Despite this initial difference in the density of active cells, there were no sex differences in the density of context cells reactivated at the time of fear recall or in the percent reactivation of any engrams established during context exposure at the time of fear recall. Together these results demonstrate that the size and reactivation of contextual engrams did not differ between the sexes but that males initially had a greater population of cells active during context exposure. One interpretation of these data is a greater role of the IL-PFC of male mice in responding to contextual information. Published literature supports this interpretation with data demonstrating significant differences between the sexes in the processing of contextual and spatial information, as well as its supporting neurobiology (28, 35, 36). Another interpretation of these data is that they demonstrate a lesser degree of habituation to Context B in male mice despite this being their second exposure to the context. As animals are repeatedly exposed to stimuli, like specific contexts, they become habituated and subsequently decrease their responsiveness, both behaviorally and cellularly (37, 38). Our results could demonstrate a reduction in habituation of IL-PFC activity in response to a familiar context specifically in male mice.

Our data showing significantly lower reactivation of IL-PFC context engrams during fear recall compared with reactivation of extinction engrams during extinction recall, regardless of sex, can be viewed through a combinatorial lens of memory recall. Fear recall in the context group consists of fear engram reactivation, which was established during fear conditioning, and contextual engram reactivation. Only the contextual engram was TRAPped and then reactivated, resulting in a relatively low percentage reactivation. In comparison, extinction recall in the extinction training group consists of fear engram reactivation, contextual engram reactivation, and extinction training engram reactivation. All three of these engrams are TRAPped during extinction training in this group, and later reactivated during recall, which is likely the cause of the greater percentage of reactivated cells. The IL-PFC is well established as a key mediator of extinction learning and recall (21) and extinction engrams have been previously identified in the mPFC (39) so the significantly greater reactivation of a putative extinction engram over a contextual engram in this region fits within our understanding of the function of the IL-PFC.

Extinction engrams, like the ones we identified in the IL-PFC, mediate successful extinction learning (9, 40, 41). The process of successful extinction learning has been shown to be new learning rather than simply the unlearning of previous fear associations (42). Fear memories persist following extinction training, but expression of fear is diminished by a relative weakening of fear memories and by competition with the newly established extinction memory (9, 40).

Heightened strength and reactivation of fear engrams has the potential of perpetuating maladaptive fear and could partially underly sex differences in fear-related learning and memory (43, 44). While our study did not directly measure engrams established during fear conditioning, these engrams are reactivated at different stages of behavioral testing and thus are captured in our analyses. In the extinction group the fear engram is initially reactivated during extinction training, and thus TRAPped. It is then subsequently reactivated, possibly to a lesser extent, during extinction recall, thus potentially also expressing C-FOS. In the context group the fear engram was not reactivated during context exposure but was reactivated on the final testing day during fear recall and thus likely expressed C-FOS. We found that females in the context group had significantly more cells in the IL-PFC that were newly activated during fear recall compared with context males. This means that a large population of cells in the IL-PFC that had not been TRAPped, potentially the fear engram, are newly activated in female mice to a greater degree than male mice. Female mice exposed to context B also showed significantly more newly activated cells in the IL-PFC (exclusively C-FOS+) upon testing for fear recall compared with female mice that underwent extinction training in context B and were being tested for extinction recall. This difference could reflect that the fear engram comes online for the first time in the female mice exposed to context B without extinction training whereas in female extinction mice the fear engram was reactivated during extinction training and thus is also TRAP+. Interestingly, males did not show this difference between context and extinction groups in newly activated cells, suggesting that this large newly activated population is a distinct feature of fear recall in females. This could indicate that in females the size of the initial fear conditioning engram is greater, an interpretation supported by studies demonstrating substantial sex differences in the recruitment of different brain regions during fear memory recall (35, 45, 46). We interpret this as potentially a greater reawakening of fear engrams in female mice during fear recall, but future work directly tagging engrams established during fear conditioning would be required to confirm this hypothesis.

To gain a more complete appreciation of potential sex differences in memory engrams it will be important to understand how both the formation and location of engrams is different between the sexes. Of the cells active during learning events, only a subset will be recruited to form a memory engram. This is determined by intrinsic properties of the cell including excitability, such that manipulating factors like CREB can bidirectionally control the allocation of specific cells to a memory engram (2, 47, 48). Intrinsic cellular properties, including excitability and CREB, are in part regulated by sex hormones, resulting in sexual dimorphism in patterns of neuronal excitability (49–51). These cellular differences between males and females could underly different patterns of engram allocation. In turn, this could mediate the sex differences we observe, such that a large population of initially active cells during context exposure in male mice does not necessarily translate to a larger or more reactivated engram. Another important factor besides what cells are allocated to a memory engram is where these cells are located.

Our analyses focused on three brain regions important for learning and memory, but in reality engrams are spread across many brain regions and function together to support memory (39, 52). Franceschini and colleagues (2023), recently demonstrated that males and females have distinct patterns of neuronal activity in different brain regions at different stages of fear recall and that functional connectivity of these regions evolves in a sexually dimorphic fashion. Like our findings, they did not observe differences in IL-PFC activity during fear recall. However, the differences we observed were uncovered only when accounting for newly active cells independent of context engrams. Future work would benefit from combining a brain-wide analyses of regions and functional connectivity with behavioral protocols that allow for the dissection of different components of memory, including context. It will also be important for future work to probe sex differences both in the allocation of cells to memory engrams and the location of these engrams as these processes fundamentally shape the cellular substrate of memories and could potentially contribute to sex differences observed in learning and memory-related dysfunction associated with PTSD.

Missing from our analyses is characterization of the cells recruited into these engrams. As previously mentioned, intrinsic cellular properties, like excitability, mediate recruitment of specific neurons during engram formation. Beyond this neuronal focus, we are beginning to appreciate that engrams are also made up of glia, specifically astrocytes. Astrocytes have been shown to express immediate early genes like C-FOS and play an integral role in engram function (53, 54). The contributions of astrocytes to engrams is a nascent area of research but a recent study demonstrated that astrocyte engram cells regulate activity of neuronal engrams and chemogenetic activation of astrocyte engram cells, independent of neurons, elicits recall (53). Recent work has also demonstrated that astrocytes are activated in response to fear learning and recall, that modulating astrocyte activity can bidirectionally control expression of fear memory, and that astrocyte activity helps facilitate the maturation of engrams during remote memory (55–58). Taken together, these findings demonstrate that astrocytes are an important player in cellular processes underlying learning and memory. Many studies, including our own, make use of C-FOS, and C-FOS dependent TRAPping, as a correlate for cellular activity without consideration of non-neuronal cell types despite evidence that astrocytes are also likely captured in these analyses. Furthermore, we know that astrocytes are sexually divergent in their neurobiology and contributions to learning and memory (59–61). A better understanding of astrocytic contributions to engrams could potentially illuminate mechanisms underlying sex differences learning and memory dysfunction associated with PTSD.

In conclusion, our study, to our knowledge, demonstrates for the first time, sex differences in IL-PFC engram activity during fear recall and context exposure. Furthermore, our work emphasizes the importance of differentiating the contributions of contextual information to engrams and highlights the preferential reactivation of extinction training engrams in the IL-PFC. This work begins to address the need for a better understanding of sex differences in memory engrams and points towards outstanding questions in the field including how cells are allocated to engrams, the contributions of non-neuronal cells to engrams, and the coordination of diverse brain regions to engram function. Addressing these questions and building on this body of work presents a promising opportunity to better understand the neurobiological underpinnings of sex differences in learning and memory related dimensions of psychiatric disorders like PTSD.

## ACKNOWLEDGMENTS

We are grateful for animal husbandry and care provided by the veterinarian and staff in the Animal Care Facility at The Saban Research Institute (TSRI). BGD’s research is supported by the National Institutes of Health (Grant Nos. R01MH134873 and R56MH128427), the Department of Pediatrics at the Keck School of Medicine of the University of Southern California, the Developmental Neuroscience and Neurogenetics Program at The Saban Research Institute, and the Child and Brain Development Program of the Canadian Institute for Advanced Research. WWT and LK received funding from the TSRI Pre-doctoral Intramural Award. The authors report no biomedical financial interests or potential conflicts of interest.

## SUPPLEMENTARY DATA AND FIGURES

**Supplemental Figure 1.**
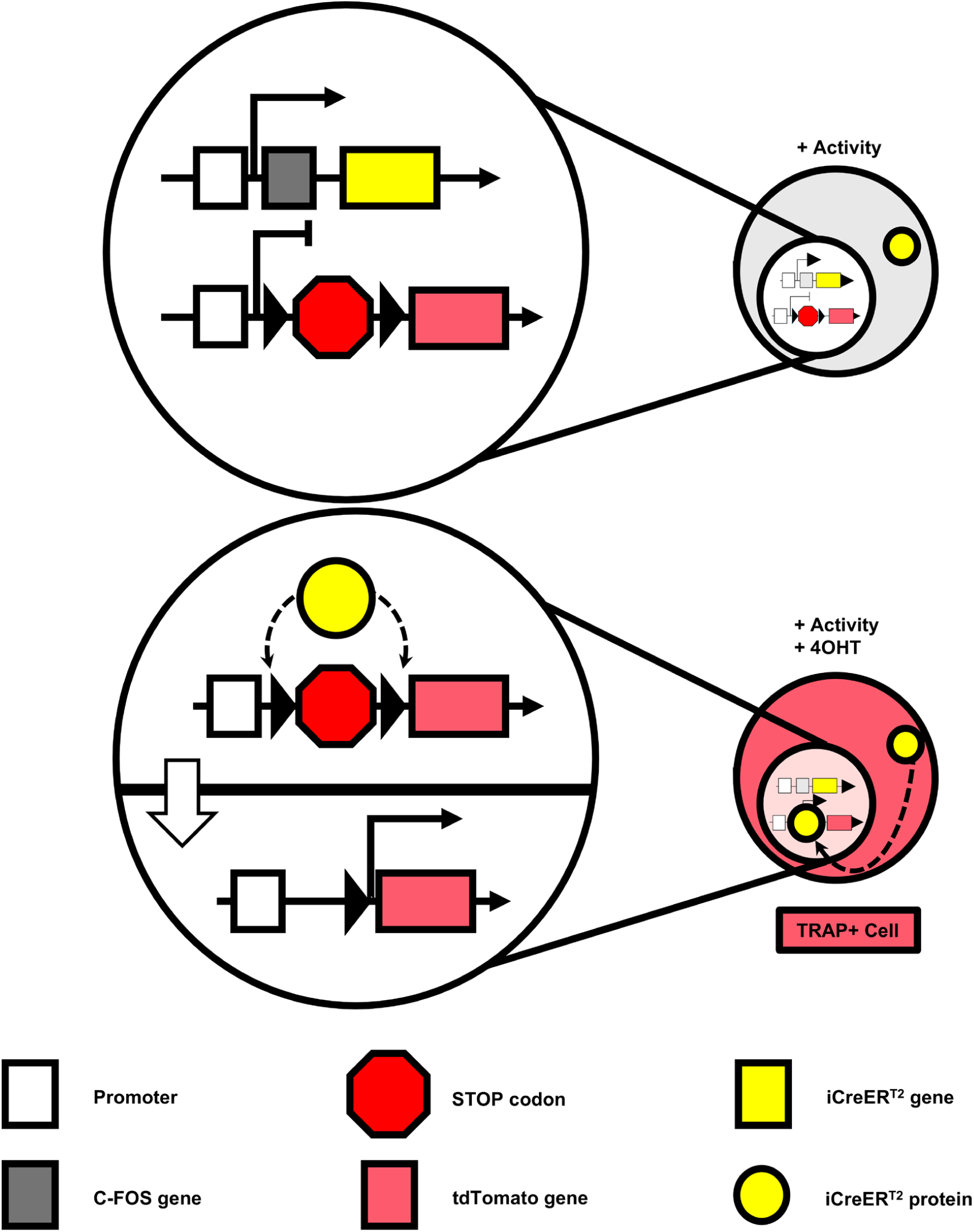
Mechanism of activity dependent cellular tagging in TRAP2;floxed-tdTomato mice. TRAP2;floxed-tdTomato express iCreER^T2^ under the C-FOS promoter, effecting the expression of iCreER^T2^ in the cytoplasm in an activity dependent manner. When 4OHT is administered, it binds iCreER^T2^ and facilitates its translocation to the nucleus. In the nucleus, iCreER^T2^ binds loxP sites flanking the STOP codon preceding the tdTomato gene. The STOP codon is excised and tdTomato is constitutively expressed in the TRAPped cell. See DeNardo et al., 2019 for details.

**Supplemental Figure 2.**
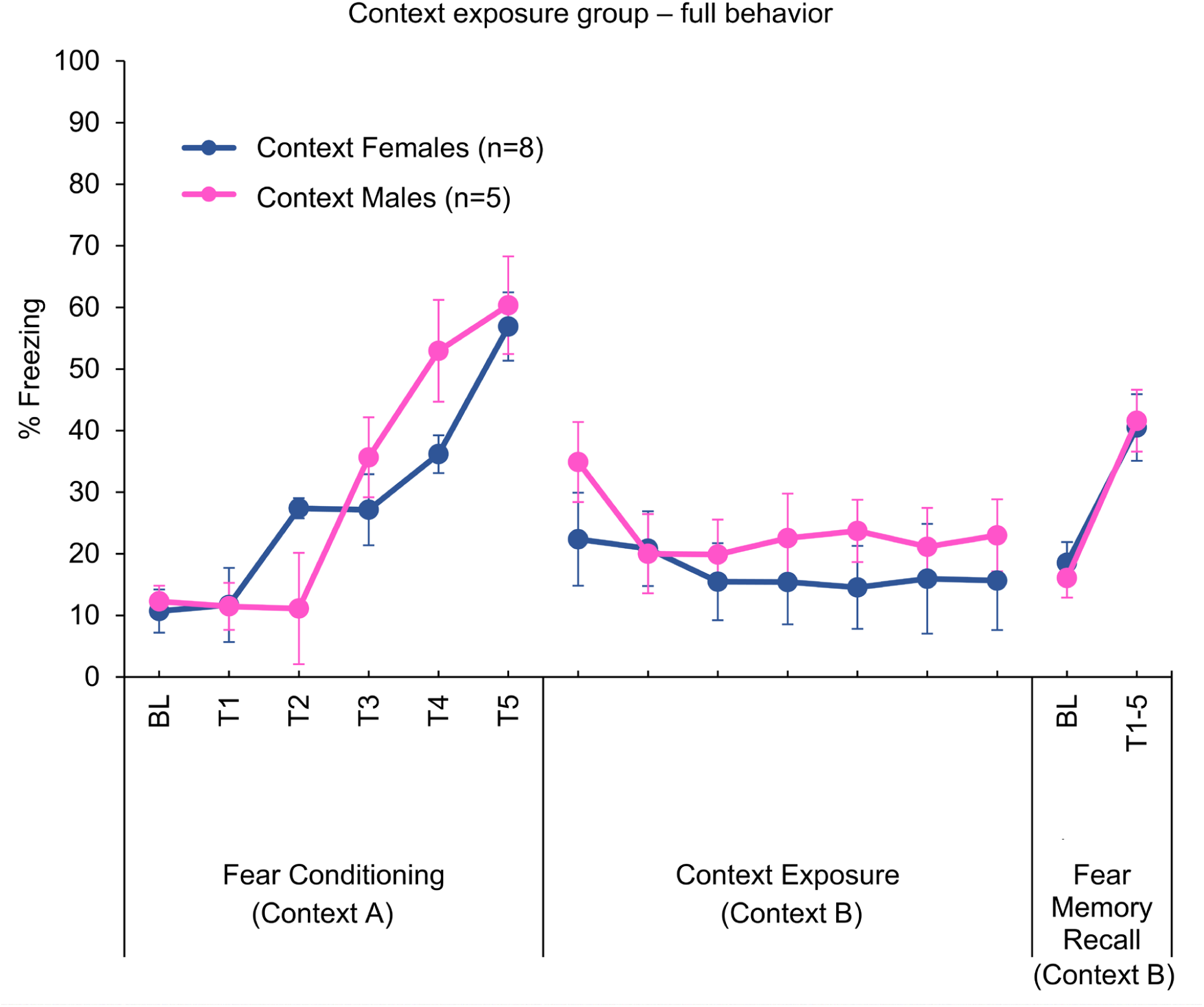
Full behavioral analyses of context exposure group during fear conditioning, contextual exposure, and fear memory recall. Both male and female mice in the context exposure group significantly increased their freezing over the course of fear conditioning in Context A. Comparing freezing during tones 1-5 by sex revealed a significant effect of tone as the mice increased their freezing (F(1, 11) = 32.785, p<0.0001), no significant effect of sex (F1, 11) = 0.088, p >0.05), and a significant interaction between tone and sex (F(1, 11) = 3.492, p = 0.02). Post-hoc testing revealed no significant differences between males and females at any specific tone presentation (all p > 0.05). Both male and female mice maintained relatively low levels of freezing during the Context B exposure (mice were not exposed to CS+ tones during context exposure and the bins demonstrated are to show freezing behavior throughout the session). There were no differences between males and females in freezing to 5 CS+ presentations during fear recall in Context B. Comparing average freezing during the full context exposure session and during the 5 tone presentations of fear recall revealed no significant effect of sex (F(1, 11) = 0.533, p > 0.05) or interaction of sex and session interaction F(1, 11) = 1.218, p > 0.05), and a significant effect of session (F(1, 11) = 33.06, p < 0.001) and post-hoc testing collapsed by sex showed that mice froze significantly more during fear recall than context exposure (V = 1, p < 0.001). Data show percent time freezing and are represented as mean±SEM.

**Supplemental Figure 3.**
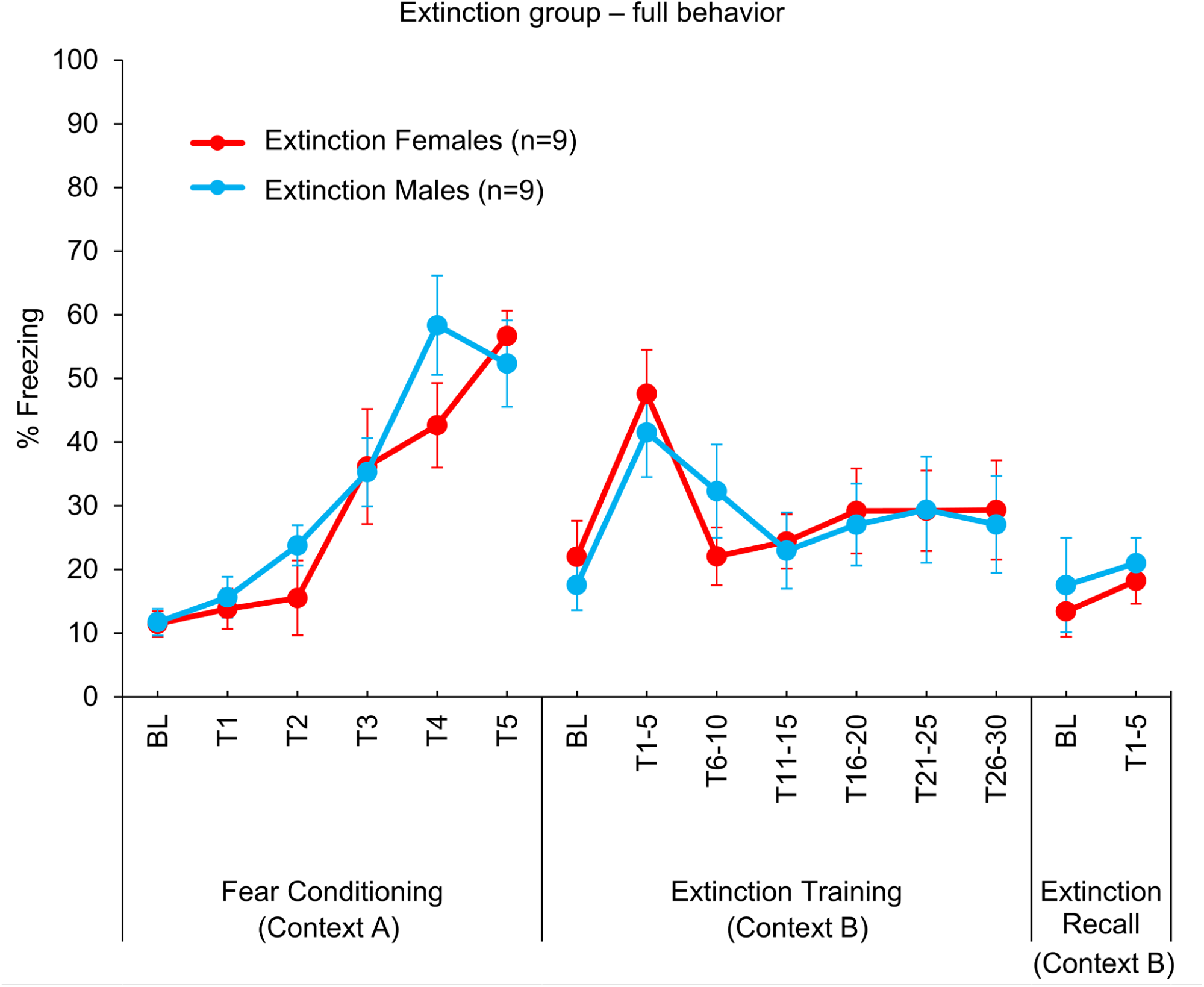
Full behavioral analyses of extinction group during fear conditioning, fear extinction training, and extinction recall. Both male and female mice in the extinction group significantly increased their freezing over the course of fear conditioning in Context A. Comparing freezing during tones 1-5 by sex revealed a significant effect of tone as the mice increased their freezing (F(1, 16) = 31.021, p < 0.0001), no significant effect of sex (F1, 16) = 0.474, p > 0.05), and no significant interaction between tone and sex (F(1, 16) = 1.549, p > 0.05). Both male and female mice decreased their freezing over the course of extinction training in Context B with no significant sex differences. Comparing average freezing during 5 tone bins by sex revealed a significant effect of bin (F(1, 16) = 8.006, p < 0.0001) and no effect of sex (F(1, 16) = 0.001, p > 0.05) or interaction between sex and bin (F(1, 16) = 1.148, p > 0.05). Both male and female mice maintained low levels of freezing during extinction recall in Context B showing no sex differences. Comparing the first 5 tone bin of extinction training with extinction recall revealed a significant main effect of testing session (F(1, 16) = 58.788, p < 0.0001), no effect of sex (F(1, 16) = 0.052, p > 0.05), and no interaction between session and sex (F(1, 16) = 1.842, p > 0.05). Posthoc testing collapsed by sex showed that mice froze significantly more during the first 5 CS+ presentations of extinction training than during extinction recall (V = 171, p < 0.0001). Data show percent time freezing and are represented as mean±SEM.

**Supplemental Figure 4.**
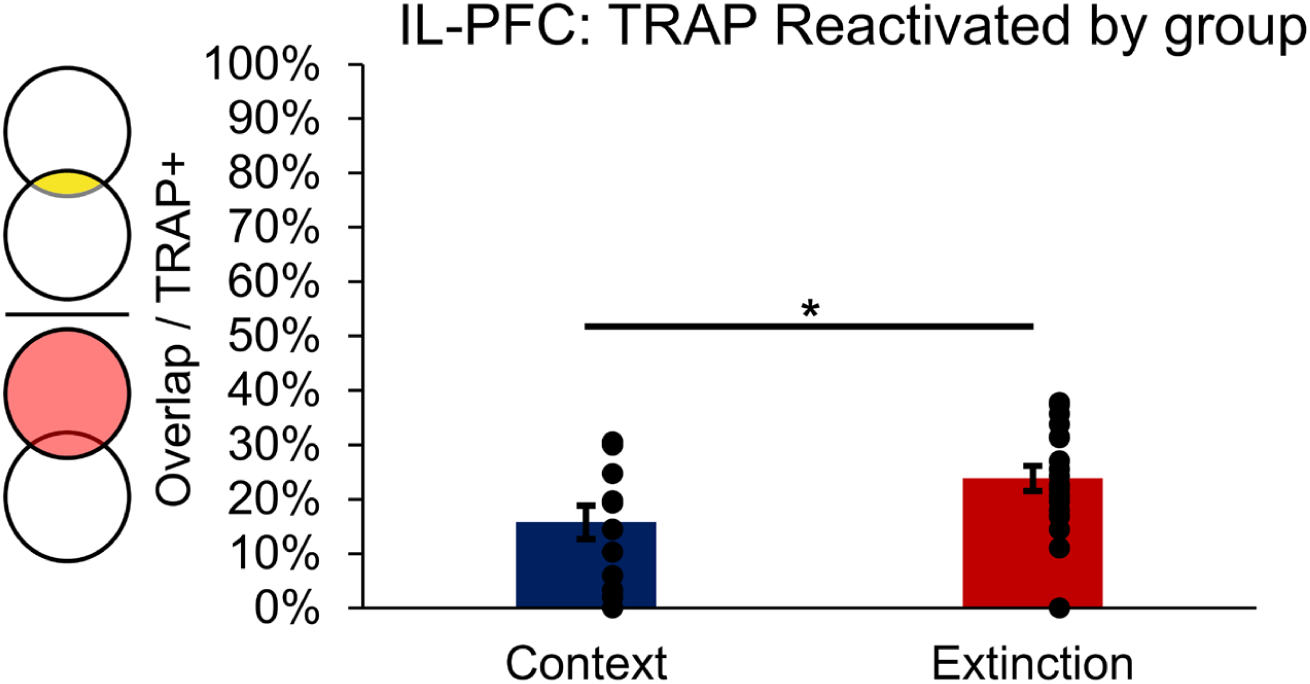
The IL-PFC is reactivated significantly more in the extinction group compared to the context group. The percentage of cells in the IL-PFC of the extinction group active during extinction training that were also active during extinction recall was greater than the percentage of cells in the context group active during context exposure that were also active during fear recall. Comparing reactivation by group and sex revealed a trending main effect of group in the IL-PFC (F(1, 27) = 3.82, p = 0.06), no main effect of sex (F(1, 27) = 2.038, p > 0.05), and no interaction between group and sex (F(1, 27) = 0.014, p < 0.05). Collapsing across sex revealed that there was significantly more reactivated cells in the extinction group than the context exposure group (W = 167.5, p = 0.045). Data show percent overlap ((TRAP+ and C-FOS+) / (all TRAP+)) and are represented as mean±SEM.

## REFERENCES.

1. Frankland PW, Josselyn SA, Kohler S (2024): Engrams. Curr Biol. 34:R559–R561.

2. Guskjolen A, Cembrowski MS (2023): Engram neurons: Encoding, consolidation, retrieval, and forgetting of memory. Mol Psychiatry. 28:3207–3219.

3. Frankland PW, Josselyn SA, Kohler S (2019): The neurobiological foundation of memory retrieval. Nat Neurosci. 22:1576–1585.

4. Josselyn SA, Tonegawa S (2020): Memory engrams: Recalling the past and imagining the future. Science. 367.

5. Han JH, Kushner SA, Yiu AP, Hsiang HL, Buch T, Waisman A, et al. (2009): Selective erasure of a fear memory. Science. 323:1492–1496.

6. Liu X, Ramirez S, Pang PT, Puryear CB, Govindarajan A, Deisseroth K, et al. (2012): Optogenetic stimulation of a hippocampal engram activates fear memory recall. Nature. 484:381–385.

7. Vetere G, Tran LM, Moberg S, Steadman PE, Restivo L, Morrison FG, et al. (2019): Memory formation in the absence of experience. Nat Neurosci. 22:933–940.

8. Tanaka KZ, Pevzner A, Hamidi AB, Nakazawa Y, Graham J, Wiltgen BJ (2014): Cortical representations are reinstated by the hippocampus during memory retrieval. Neuron. 84:347–354.

9. Lacagnina AF, Brockway ET, Crovetti CR, Shue F, McCarty MJ, Sattler KP, et al. (2019): Distinct hippocampal engrams control extinction and relapse of fear memory. Nat Neurosci. 22:753–761.

10. 10. Hsiang HL, Epp JR, van den Oever MC, Yan C, Rashid AJ, Insel N, et al. (2014): Manipulating a “cocaine engram” in mice. J Neurosci. 34:14115–14127.

11. Cowansage KK, Shuman T, Dillingham BC, Chang A, Golshani P, Mayford M (2014): Direct reactivation of a coherent neocortical memory of context. Neuron. 84:432–441.

12. Redondo RL, Kim J, Arons AL, Ramirez S, Liu X, Tonegawa S (2014): Bidirectional switch of the valence associated with a hippocampal contextual memory engram. Nature. 513:426–430.

13. Velasco ER, Florido A, Milad MR, Andero R (2019): Sex differences in fear extinction. Neurosci Biobehav Rev. 103:81–108.

14. Lebron-Milad K, Milad MR (2012): Sex differences, gonadal hormones and the fear extinction network: implications for anxiety disorders. Biol Mood Anxiety Disord. 2:3.

15. Ramikie TS, Ressler KJ (2018): Mechanisms of Sex Differences in Fear and Posttraumatic Stress Disorder. Biol Psychiatry. 83:876–885.

16. DeNardo LA, Liu CD, Allen WE, Adams EL, Friedmann D, Fu L, et al. (2019): Temporal evolution of cortical ensembles promoting remote memory retrieval. Nat Neurosci. 22:460–469.

17. Bankhead P, Loughrey MB, Fernandez JA, Dombrowski Y, McArt DG, Dunne PD, et al. (2017): QuPath: Open source software for digital pathology image analysis. Sci Rep. 7:16878.

18. Venkataraman A, Dias BG (2023): Expanding the canon: An inclusive neurobiology of thalamic and subthalamic fear circuits. Neuropharmacology. 226:109380.

19. Venkataraman A, Hunter SC, Dhinojwala M, Ghebrezadik D, Guo J, Inoue K, et al. (2021): Incerto-thalamic modulation of fear via GABA and dopamine. Neuropsychopharmacology. 46:1658–1668.

20. Venkataraman A, Brody N, Reddi P, Guo J, Gordon Rainnie D, Dias BG (2019): Modulation of fear generalization by the zona incerta. Proc Natl Acad Sci U S A. 116:9072–9077.

21. Giustino TF, Maren S (2015): The Role of the Medial Prefrontal Cortex in the Conditioning and Extinction of Fear. Front Behav Neurosci. 9:298.

22. Sotres-Bayon F, Quirk GJ (2010): Prefrontal control of fear: more than just extinction. Curr Opin Neurobiol. 20:231–235.

23. Orsini CA, Maren S (2012): Neural and cellular mechanisms of fear and extinction memory formation. Neurosci Biobehav Rev. 36:1773–1802.

24. Chou XL, Wang X, Zhang ZG, Shen L, Zingg B, Huang J, et al. (2018): Inhibitory gain modulation of defense behaviors by zona incerta. Nat Commun. 9:1151.

25. Zhao P, Zhao M, Wang H, Jiang T, Jia X, Tian J, et al. (2020): Long-range inputome of cortical neurons containing corticotropin-releasing hormone. Sci Rep. 10:12209.

26. Milad MR, Zeidan MA, Contero A, Pitman RK, Klibanski A, Rauch SL, et al. (2010): The influence of gonadal hormones on conditioned fear extinction in healthy humans. Neuroscience. 168:652–658.

27. Milad MR, Pitman RK, Ellis CB, Gold AL, Shin LM, Lasko NB, et al. (2009): Neurobiological basis of failure to recall extinction memory in posttraumatic stress disorder. Biol Psychiatry. 66:1075–1082.

28. Fleischer AW, Frick KM (2023): New perspectives on sex differences in learning and memory. Trends Endocrinol Metab. 34:526–538.

29. 29. Gall CM, Le AA, Lynch G (2023): Sex differences in synaptic plasticity underlying learning. J Neurosci Res. 101:764–782.

30. Franceschini A, Mazzamuto G, Checcucci C, Chicchi L, Fanelli D, Costantini I, et al. (2023): Brain-wide neuron quantification toolkit reveals strong sexual dimorphism in the evolution of fear memory. Cell Rep. 42:112908.

31. Mashour GA, Pal D, Brown EN (2022): Prefrontal cortex as a key node in arousal circuitry. Trends Neurosci. 45:722–732.

32. Zhang Q, Weber MA, Narayanan NS (2021): Medial prefrontal cortex and the temporal control of action. Int Rev Neurobiol. 158:421–441.

33. Mair RG, Francoeur MJ, Krell EM, Gibson BM (2022): Where Actions Meet Outcomes: Medial Prefrontal Cortex, Central Thalamus, and the Basal Ganglia. Front Behav Neurosci. 16:928610.

34. Hockley A, Malmierca MS (2024): Auditory processing control by the medial prefrontal cortex: A review of the rodent functional organisation. Hear Res. 443:108954.

35. Keiser AA, Turnbull LM, Darian MA, Feldman DE, Song I, Tronson NC (2017): Sex Differences in Context Fear Generalization and Recruitment of Hippocampus and Amygdala during Retrieval. Neuropsychopharmacology. 42:397–407.

36. Yagi S, Lee A, Truter N, Galea LAM (2022): Sex differences in contextual pattern separation, neurogenesis, and functional connectivity within the limbic system. Biol Sex Differ. 13:42.

37. Merchie A, Gomot M (2023): Habituation, Adaptation and Prediction Processes in Neurodevelopmental Disorders: A Comprehensive Review. Brain Sci. 13.

38. Wilson DA, Linster C (2008): Neurobiology of a simple memory. J Neurophysiol. 100:2–7.

39. Gu X, Wu YJ, Zhang Z, Zhu JJ, Wu XR, Wang Q, et al. (2022): Dynamic tripartite construct of interregional engram circuits underlies forgetting of extinction memory. Mol Psychiatry. 27:4077–4091.

40. Zhang X, Kim J, Tonegawa S (2020): Amygdala Reward Neurons Form and Store Fear Extinction Memory. Neuron. 105:1077–1093 e1077.

41. Lee H, Kaang BK (2023): How engram mediates learning, extinction, and relapse. Curr Opin Neurobiol. 81:102723.

42. Dunsmoor JE, Niv Y, Daw N, Phelps EA (2015): Rethinking Extinction. Neuron. 88:47–63.

43. Inslicht SS, Metzler TJ, Garcia NM, Pineles SL, Milad MR, Orr SP, et al. (2013): Sex differences in fear conditioning in posttraumatic stress disorder. J Psychiatr Res. 47:64–71.

44. Day HLL, Stevenson CW (2020): The neurobiological basis of sex differences in learned fear and its inhibition. Eur J Neurosci. 52:2466–2486.

45. Bauer EP (2023): Sex differences in fear responses: Neural circuits. Neuropharmacology. 222:109298.

46. Reppucci CJ, Petrovich GD (2018): Neural substrates of fear-induced hypophagia in male and female rats. Brain Struct Funct. 223:2925–2947.

47. Kim J, Kwon JT, Kim HS, Josselyn SA, Han JH (2014): Memory recall and modifications by activating neurons with elevated CREB. Nat Neurosci. 17:65–72.

48. Yiu AP, Mercaldo V, Yan C, Richards B, Rashid AJ, Hsiang HL, et al. (2014): Neurons are recruited to a memory trace based on relative neuronal excitability immediately before training. Neuron. 83:722–735.

49. Fabian CB, Seney ML, Joffe ME (2023): Sex differences and hormonal regulation of metabotropic glutamate receptor synaptic plasticity. Int Rev Neurobiol. 168:311–347.

50. Brann DW, Lu Y, Wang J, Sareddy GR, Pratap UP, Zhang Q, et al. (2021): Neuron-Derived Estrogen-A Key Neuromodulator in Synaptic Function and Memory. Int J Mol Sci. 22.

51. 51. Guo G, Kang L, Geng D, Han S, Li S, Du J, et al. (2020): Testosterone modulates structural synaptic plasticity of primary cultured hippocampal neurons through ERK - CREB signalling pathways. Mol Cell Endocrinol. 503:110671.

52. Roy DS, Park YG, Kim ME, Zhang Y, Ogawa SK, DiNapoli N, et al. (2022): Brain-wide mapping reveals that engrams for a single memory are distributed across multiple brain regions. Nat Commun. 13:1799.

53. Williamson MR, Kwon W, Woo J, Ko Y, Maleki E, Yu K, et al. (2025): Learning-associated astrocyte ensembles regulate memory recall. Nature. 637:478–486.

54. Cruz-Mendoza F, Jauregui-Huerta F, Aguilar-Delgadillo A, Garcia-Estrada J, Luquin S (2022): Immediate Early Gene c-fos in the Brain: Focus on Glial Cells. Brain Sci. 12.

55. Suthard RL, Senne RA, Buzharsky MD, Pyo AY, Dorst KE, Diep AH, et al. (2023): Basolateral Amygdala Astrocytes Are Engaged by the Acquisition and Expression of a Contextual Fear Memory. J Neurosci. 43:4997–5013.

56. Suthard RL, Senne RA, Buzharsky MD, Diep AH, Pyo AY, Ramirez S (2024): Engram reactivation mimics cellular signatures of fear. Cell Rep. 43:113850.

57. Refaeli R, Kreisel T, Yaish TR, Groysman M, Goshen I (2024): Astrocytes control recent and remote memory strength by affecting the recruitment of the CA1-->ACC projection to engrams. Cell Rep. 43:113943.

58. Yamao H, Matsui K (2025): Astrocytic determinant of the fate of long-term memory. Glia. 73:309–329.

59. Gozlan E, Lewit-Cohen Y, Frenkel D (2024): Sex Differences in Astrocyte Activity. Cells. 13.

60. Meadows SM, Palaguachi F, Jang MW, Licht-Murava A, Barnett D, Zimmer TS, et al. (2024): Hippocampal astrocytes induce sex-dimorphic effects on memory. Cell Rep. 43:114278.

61. Taylor WW, Imhoff BR, Sathi ZS, Liu WY, Garza KM, Dias BG (2021): Contributions of glucocorticoid receptors in cortical astrocytes to memory recall. Learn Mem. 28:126–133.

